# An intrinsic temporal order of c-Jun N-terminal phosphorylation regulates its activity by orchestrating co-factor recruitment

**DOI:** 10.1101/2021.11.03.465096

**Authors:** Christopher A. Waudby, Saul Alvarez-Teijeiro, Simon Suppinger, Paul R. Brown, Axel Behrens, John Christodoulou, Anastasia Mylona

**Affiliations:** Institute of Structural and Molecular Biology, Birkbeck College, London; Institute of Structural and Molecular Biology, University College London, U.K.; Randall Division of Cell and Molecular Biophysics, Guy’s Campus, King’s College, London, U.K.; Cancer Stem Cell Laboratory, Institute of Cancer Research, London, U.K.; Imperial College, Division of Cancer, Department of Surgery and Cancer; Convergence Science Centre, Imperial College, London, SW7 2BU, U.K.

## Abstract

Protein phosphorylation is a major regulatory mechanism of cellular signalling. The c-Jun proto-oncoprotein is phosphorylated at four residues within its transactivation domain (TAD) by the JNK family kinases, but the functional significance of c-Jun multisite phosphorylation has remained elusive. Here we show that c-Jun phosphorylation by JNK exhibits a defined temporal kinetics, with serine63 and serine73 being phosphorylated more rapidly than threonine91 and threonine93. We identified the positioning of the phosphorylation sites relative to the kinase docking motif, and their primary sequence, as the main factors controlling phosphorylation kinetics. Functional analysis revealed three c-Jun phosphorylation states: unphosphorylated c-Jun recruits the Mbd3 repressor, serine63/73 doubly-phosphorylated c-Jun binds to the Tcf4 co-activator, whereas the fully phosphorylated form disfavours Tcf4 binding attenuating JNK signalling. Thus, c-Jun phosphorylation encodes multiple functional states that drive a complex signalling response from a single JNK input.

## Introduction

Protein phosphorylation is the most abundant and important post-translational protein modification^1,2^ and is a major regulatory mechanism of cellular signalling^3^. Deregulation of phosphorylation pathways commonly underlies disease aetiology^4^ Proteins often are phosphorylated not only at one but at multiple sites by a single kinase. Multisite phosphorylation is a key regulatory mechanism in signalling that controls many cellular processes, and has been suggested to provide a precise tool to generate complex functional outputs, often from a single kinase input^5–9^. This is often achieved via interactions between the multisite phosphorylated protein and partner proteins, and these phosphorylation-dependent interactions determine the biological output^10^. Transcription is an important biological process regulated by multisite phosphorylation^11,12^. We have previously used a pipeline that facilitated a mechanistic understanding of the temporal dynamics of multisite phosphorylation events by the ERK kinase in the case of the Ternary Complex Factor (TCF) Elk-1, transcriptional coactivator^13^. Phosphorylation at eight S/T-P (Ser or Thr-Pro) sites occurred with a defined kinetics, leading to intrinsic temporal phosphorylation events that effectively trigger and subsequently limit its transcriptional response by promoting and then inhibiting the recruitment of the Mediator complex^13^. Thus, understanding the intrinsic kinetics of multisite phosphorylation is key to gain insights on how collectively multiple phosphorylated sites by a kinase within a domain can shape the transcriptional response.

AP-1 (activator protein 1) activity is strongly induced in response to numerous signals including growth factors, cytokines and extracellular stress^14^ AP-1 consists of heterodimeric complexes formed by members of the Jun family (c-Jun, JunB and JunD) together with the Fos proteins (c-Fos, FosB, Fra1 and Fra2) and members of the ATF and CREB families. The protooncoprotein c-Jun is a major component of the AP-1 transcription factor that controls the expression of many genes important for cell proliferation and tumorigenesis^15,16^.

An important mechanism to stimulate AP-1 function is the N-terminal phosphorylation of the c-Jun transactivation domain (TAD) at multiple S/T-P sites by the c-Jun N-terminal kinase (JNK) group of mitogen-activated protein kinases (MAPK)^17^ The JNKs have three isoforms: JNK1, JNK2 and JNK3. JNK1 and JNK2 are expressed in almost all tissues, whereas the expression of JNK3 is restricted to neuronal tissues. Both JNK1 and JNK2 actively contribute to c-Jun phosphorylation, and are at least partly redundant^17^

JNKs phosphorylate the c-Jun TAD at four S/T-P residues, S63, S73, T91 and T93 (Fig. 1A), which modulates its interaction with various partner proteins to control target gene transcription^16,18–20^. The unphosphorylated c-Jun TAD interacts with Mbd3, a subunit of the nucleosome remodelling and histone deacetylation (NuRD) repressor complex, which in turn recruits NuRD to AP-1 target genes to mediate gene repression. This repression is relieved by c-Jun phosphorylation following JNK activation^21^. Phosphorylated c-Jun at its TAD by JNK interacts with the Tcf4 HMG-box transcription factor activating the *c-jun* promoter^16^.

**Figure 1.**
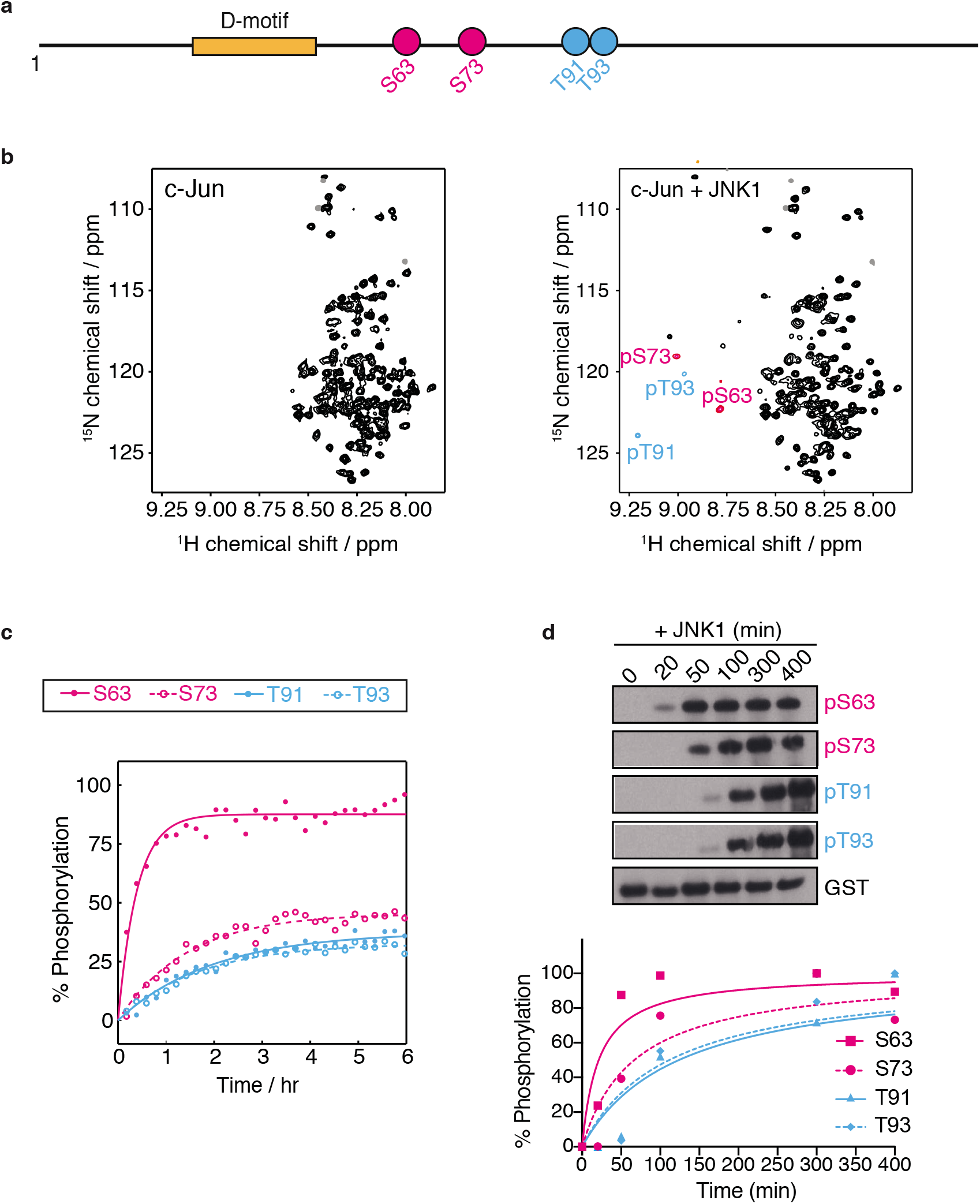
Multisite phosphorylation kinetics of the c-Jun TAD by JNK1. (**a**) Schematic representation of the c-Jun TAD sequence, showing the location of the S/T-P phosphorylation sites and the JNK binding motif (D-motif). (**b**) 2D^1^H,^15^N correlation spectra of (left) unphosphorylated c-Jun TAD, and (right) following 6 hr *in vitro* phosphorylation by JNK1. Resonances from phosphorylated residues are highlighted with pink or cyan colouring corresponding to fast or slow sites respectively, with assignments as indicated. (**c**) Site-specific kinetics of c-Jun phosphorylation by JNK1 obtained from time-resolved NMR measurements. Data are fitted to single exponential build-ups. (**d**) Western blot analysis of the *in vitro* phosphorylation kinetics of recombinant c-Jun TAD by JNK1 using phosphorylation-specific antibodies (top) and quantification of the detected protein levels by western blot using Image Studio Lite Software (Licor) and normalized to total GST c-Jun TAD (bottom).

The multisite N-terminal phosphorylation of c-Jun by JNK is also a key event linked to tumorigenesis. Mice in which the c-Jun phosphorylation sites S63 and S73 are mutated to alanines *(JunAA/JunAA)* develop normally but show specific defects in oncogenic transformation^22^. Moreover, in the intestine both Mbd3 and Tcf4/*b*-catenin control cellular proliferation and tumorigenesis via JNK/c-Jun N-terminal phosphorylation. Mice lacking Mbd3 (*mbd3*^ΔG/ΔG^) show increased intestinal progenitor proliferation, which can be rescued upon deletion of one *c-jun* allele (*mbd3*^ΔG/ΔG^; *c-jun*^ΔG/+^)^21^. In a mouse model of intestinal cancer that is heterozygous for a nonsense mutation at codon 850 of the *Apc* gene (*Apc*^Min/+^), mice that are crossed to *JunAA* mice (*Apc*^Min/+^; *JunAA/JunAA*) exhibit greatly reduced intestinal tumors comparing to the *Apc*^Min/+^ mice^16^. However, although the biological significance of JNK c-Jun S63 and S73 N-terminal phosphorylation and its role in tumour development have been clearly demonstrated, the roles of the individual four phosphorylation sites and the interplay among them, and particularly how they might cooperate to regulate the temporal cofactor recruitment and thus transcriptional output, have not yet been explored.

In this study, we show that c-Jun N-terminal multisite phosphorylation by JNK occurs with a range of rates, leading to intrinsic temporal phosphorylation states, both *in vitro* and *in vivo.* This differential kinetic behaviour arises from differences in the primary sequence of the phosphorylation sites, as well as their position relative to the kinase docking motif. Finally, we examine the functional impact of the differential kinetics of c-Jun N-terminal phosphorylation and identify that c-Jun TAD exhibits three principal temporal phosphorylation states: an unphosphorylated repressive state that binds the Mbd3 repressor, a double phosphorylated transcriptionally active state that binds the Tcf4 transcriptional activator, and finally a fully phosphorylated inactive form that disfavours Tcf4 binding and thus attenuates JNK signalling. Our results, therefore, demonstrate that the c-Jun N-terminal phosphorylation is a sequencial phosphorylayion cascade that controls in a time-dependent manner recruitment of c-Jun cofactors, threreby auto adjusting its transcriptional output via a single JNK kinase signalling input.

## Results

### Differential kinetics of the N-terminal multisite phosphorylation of c-Jun by JNK

As a first step to explore the phosphorylation kinetics of the c-Jun TAD by JNK, we used time-resolved NMR spectroscopy to monitor the modification of individual c-Jun phosphosites in real time^23–25^. We expressed and purified c-Jun TAD (residues 1-151) containing all four phosphorylation sites, S63, S73, T91 and T91 and the MAPK binding motif (D-motif, residues 32-50) which controls phosphorylation of these sites^26,27^ (Fig. 1a). The narrow chemical shift dispersion in the 2D^1^H,^15^N correlation spectrum of unphosphorylated c-Jun TAD (Fig. 1b) indicates that the domain is intrinsically disordered, and following backbone resonance assignment (Fig. S1a) an analysis of secondary chemical shifts^28^ showed no significant populations of secondary structure (Fig. S1b), consistent with previous reports of c-Jun fragments comprising residues 1-123 and 1-276^29^.

We reacted^15^N-labelled c-Jun TAD with recombinant active JNK1 and performed time-resolved NMR to monitor its modification rate *in vitro* (Fig. 1b & S2). Due to the change in their chemical environment, phosphorylated serine and threonine (pS and pT) amide resonances appear in a distinctive and well-resolved region of the 2D^1^H,^15^N correlation spectrum^30^, and their intensities can therefore be readily tracked as a function of time (Fig. 1c). pS and pT resonances that were observed to build up over time were assigned to the four phosphorylation sites via sequential site-directed mutagenesis to alanine (i.e. S63A, S73A, T91A and T93A) to eliminate phosphorylation (Fig. S3a). The narrow chemical shift dispersion observed throughout the phosphorylation reaction (Fig. 1b and S2) indicates that the c-Jun TAD domain remains disordered in all its phosphorylation states, consistent with previous analyses of hyperphosphorylated forms of c-Jun 1-123 and 1-276 by ERK2^29^.

Following addition of JNK1, the build-up of phosphorylated resonances displayed a range of rates, in the order S63 > S73 > T91 ≈ T93 (Fig. 1c). Phosphorylation of all residues followed exponential kinetics with no discernible lag phases, which provided early evidence that there is no strong cooperativity between adjacent phosphosites. This is explored in further detail below. The kinetics observed by NMR were confirmed by immunoblotting of the *in vitro* reactions with phosphorylation-specific antibodies against each of the phosphorylated residues (Fig. 1d). The site specificity of the these phospho-antibodies was verified using alanine mutants for each phosphorylation site (Fig. S3b).

To examine the N-terminal phosphorylation kinetics of c-Jun *in vivo*, we used anisomycin treatment of NIH 3T3 cells to stimulate JNK phosphorylation and activation. This occurred with a half-time of about 12 min following a ca. 5 min lag period, and then remained in its active phosphorylated form for the 30 min duration of measurements (Fig. 2a). We then monitored the phosphorylation rates of S63, S73, T91 and T93 within endogenous c-Jun by immunoblotting (Fig. 2b). Consistent with our *in vitro* measurements, phosphorylation of S63 and S73 occurred rapidly following JNK activation, with half-times of ca. 12 min, while phosphorylation of T91 and T93 occurred more slowly, with a half-time of ca. 15-17 min (Fig. 2b). Taken together, these data indicate that phosphorylation of the c-Jun TAD proceeds under physiological conditions through two phases, in which S63 and S73 are phosphorylated more rapidly followed by T91 and T93 (Fig. 2c).

**Figure 2.**
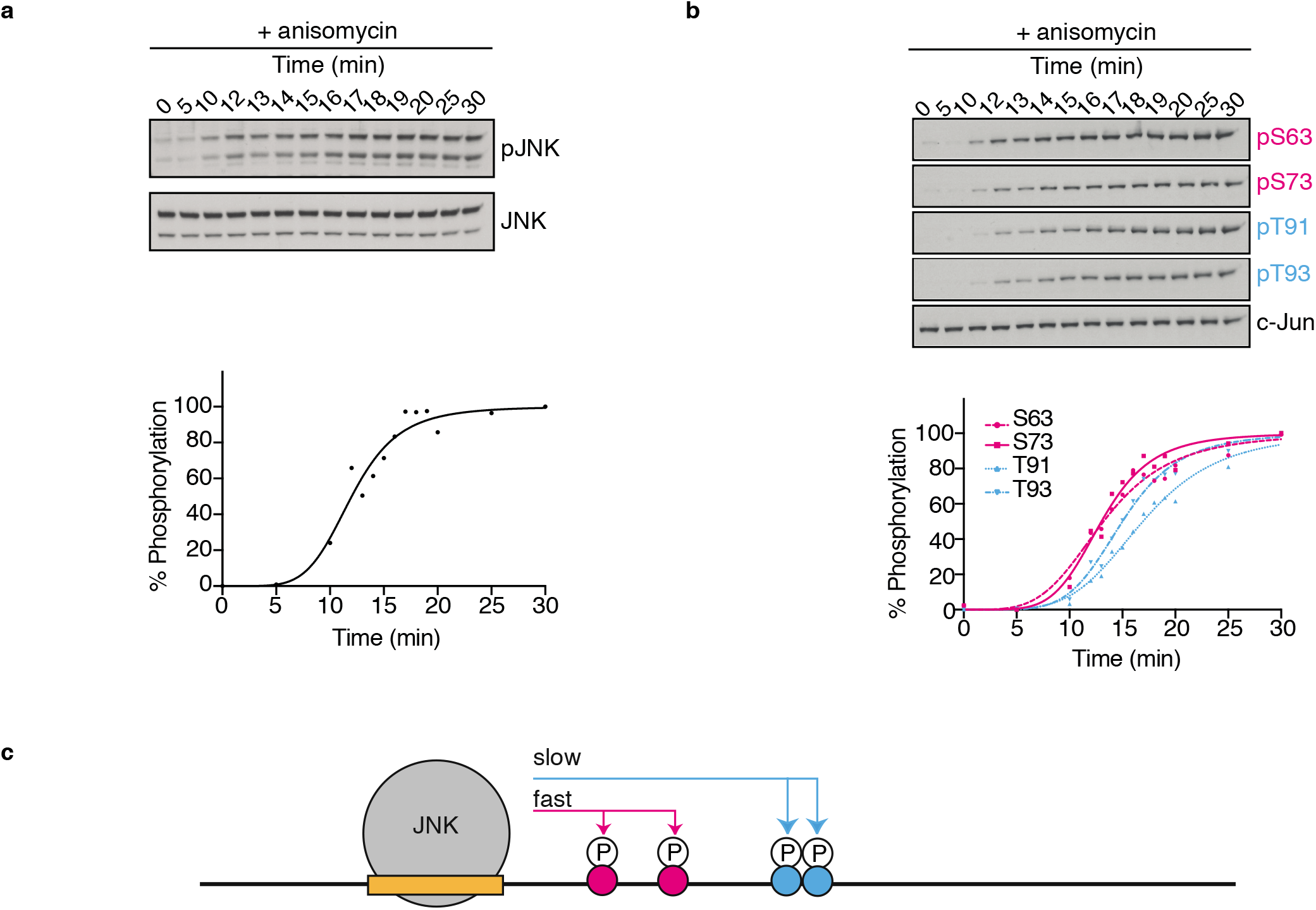
Kinetics of the c-Jun N-terminal phosphorylation *in vivo* (**a**) Western blot analysis of the phosphorylation (activation) of endogenous JNK in NIH3T3 cells following anisomycin treatment (top), and quantification of the detected protein levels using Image Studio Lite Software (Licor) and normalized to total JNK (bottom). (**b**) Western blot analysis of the phosphorylation kinetics of endogenous c-Jun in NIH3T3 cells following anisomycin treatment using phosphorylation-specific antibodies (top), and quantification of the detected protein levels using Image Studio Lite Software (Licor) and normalized to total c-Jun (bottom). (**c**) Schematic summary of the observed phosphorylation kinetics of the c-Jun N-terminal phosphorylation by JNK.

### JNK1 kinase operates independently on c-Jun phosphorylation sites

Having identified variations in the phosphorylation rates of c-Jun TAD residues, we sought to understand their molecular basis. To do this, we first examined whether any cooperativity existed in the phosphorylation of different residues. As noted above, the phosphorylation kinetics of individual residues could be fitted closely by exponential functions, without any evidence of lag phases (Fig. 1c), which indicated that phosphorylation of a particular site was not a pre-requisite for phosphorylation of another site. This conclusion is further supported by analysis of the phosphorylation kinetics of the single-site alanine variants, S63A, S73A, T91A and T93A, using time-resolved NMR. Alanine replacement of any of the individual sites did not prevent efficient phosphorylation, at equivalent rates to that of the wildtype protein, of any of the subsequent phosphorylated sites (Fig. 3a, S4a). To the contrary, the phosphorylation of T93 within the T91A variant was ca. 3 times more rapid than in the wildtype (Fig. 3a and S4a). Given the proximity of these residues, we suggest that this is most likely an effect of the local sequence on the catalytic activity of JNK1.

**Figure 3.**
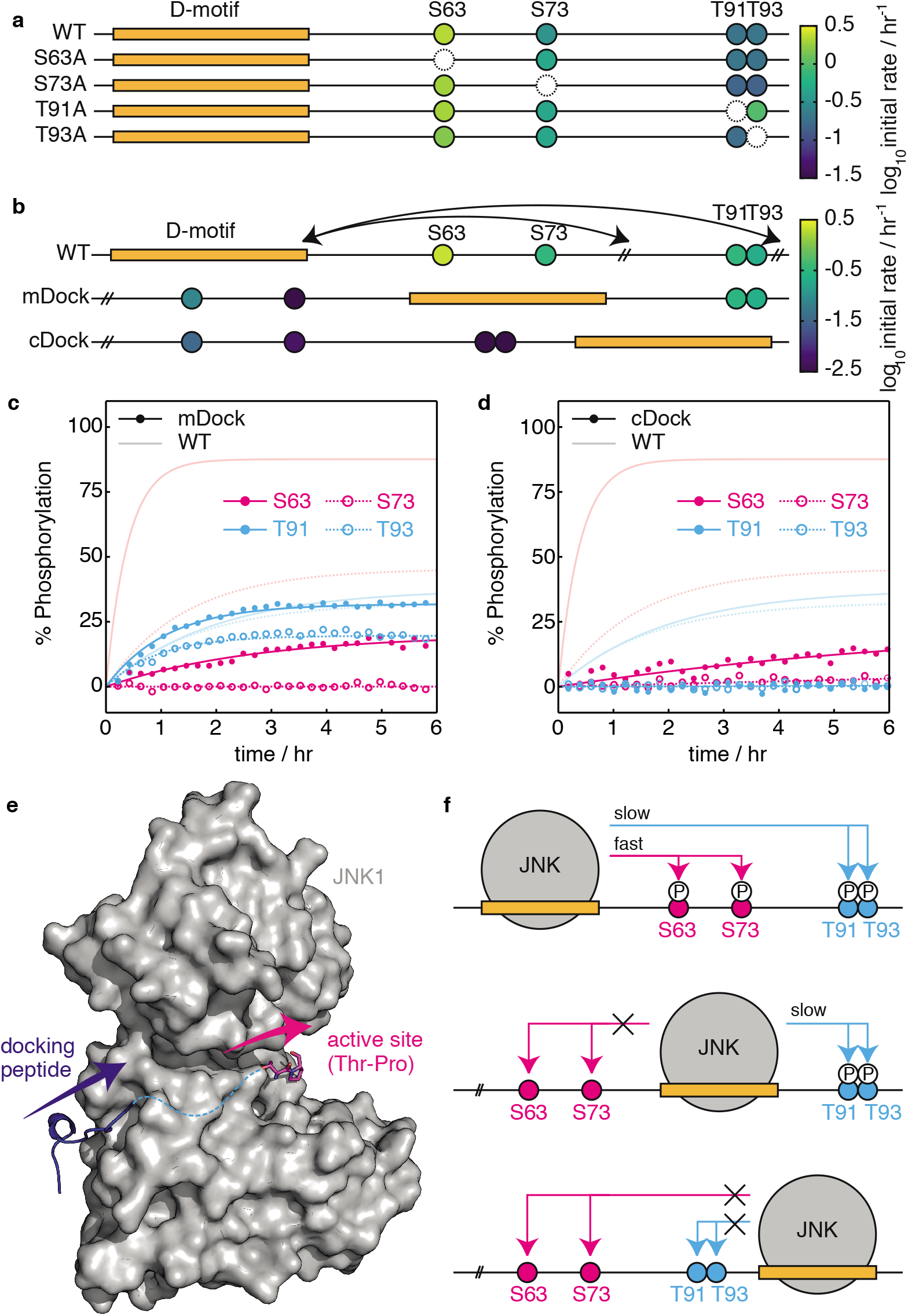
Effect of relative distance and orientation of the S/T-P sites from the D-motif on the c-Jun N-terminal progressive multisite phosphorylation. (**a**) Schematic representation of the c-Jun TAD constructs S63A, S73A, T91A and T93A. The initial rate of phosphorylation of residues by JNK1, determined by time-resolved NMR spectroscopy, is indicated according to the displayed colour scale. (**b**) Schematic representation of the c-Jun mDock and cDock constructs comparing to wild-type (WT). Initial phosphorylation rates measured using time-resolved NMR spectroscopy are indicated according to the displayed colour scale. (**c, d**) Sitespecific buildup curves of the phosphorylation of c-Jun (**c**) mDock and (**d**) cDock variants by JNK1; blurred lines show wild-type c-Jun TAD for comparison. (**e**) Crystal structure of JNK1 (grey surface) bound to a D-motif peptide (dark blue), superimposed with a crystal structure of a consensus T-P substract peptide (magenta) bound to the homologous DYRK kinase (not shown). Arrows indicate the orientation of the bound peptides, with a hypothetical linker sequence indicated (dashed cyan). (**f**) Schematic representation summarizing the effect of the relative distance and orientation of the c-Jun TAD S/T-P residues from the D-motif on the temporal dynamics of the c-Jun N-terminal phosphorylation.

### Efficiency of c-Jun N-terminal phosphorylation by JNK1 is determined by the orientation of the phosphorylation sites relative to the D-motif

To further understand the factors underlying the variation in c-Jun phosphorylation rates by JNK1, we hypothesised that differences in kinetics may reflect the varying spatial proximity of phosphorylation sites relative to the D-motif, particularly given that the fast S63 and S73 sites are located closer to the D-motif than the slower T91 and T93 sites. To test this, we engineered two c-Jun TAD constructs, mDock and cDock, in which the position of the D-motif was transposed relative to the fast and slow groups of phosphorylation sites (Fig. 3b and S4b).

We measured the phosphorylation kinetics of mDock and cDock variants using time-resolved NMR (Fig. 3b,c,d & S3a). The increased proximity of T91 and T93 in the mDock variant did not result in more rapid phosphorylation, but instead their phosphorylation rates remained very similar to those in the wild-type TAD construct (Fig. 3c). However, the phosphorylation of sites N-terminal to the D-motif was strongly suppressed, with only very slow modification of the normally fast S63 site observed in both mDock and cDock variants (Fig. 3c,d). These observations can be rationalised by consideration of the three-dimensional structural model of JNK1, showing a bound D-motif peptide^31^ and a consensus T-P substrate peptide (generated by alignment of a crystal structure of DYRK)^32^ (Fig. 3e). These peptides bind to the enzyme in a common orientation, with only a short distance from the C-terminal of the D-motif to the N-terminal of the phosphorylation site, such that phosphorylation sites C-terminal to the D-motif can readily access the active site. In contrast, phosphorylation sites N-terminal to the D-motif require the intervening linker (if long enough) to fold back upon itself at a high entropic cost, resulting in a significantly slower rate of phosphorylation. Taken together, these data indicate that although the relative distance of the phosphorylation sites to the D-motif is not essential in defining the temporal order of c-Jun multisite phosphorylation by JNK1, efficient phosphorylation does require positioning of the phosphorylation sites downstream of the D-motif (Fig. 3f).

### Primary sequence strongly governs the rate of c-Jun N-terminal phosphorylation by JNK1

Next, we sought to differentiate the effects of the primary sequence of c-Jun and the distance of phospho-sites from the D-motif on the rate of phosphorylation by JNK1. For this purpose, we engineered a ‘swap’ variant, in which a sequence containing the fast S63 and S73 phospho-sites was exchanged with a sequence containing the slower T91 and T93 sites (Fig. 4a & S4c), and again measured the phosphorylation rates of individual sites by time-resolved NMR (Fig. 4a, b & S3a). We observed that the rate of S63 phosphorylation decreased ca. 4-fold, which may be consistent with a decrease in its effective concentration given the longer linker length^33^. However, both residues, despite now being further away from the D-motif than T91 and T93, were still phosphorylated with higher rates than T91 and T93, whose modification rate remained at similar levels than the wild-type (Fig. 4a, b). This is further evidenced by the fact that in the 2SA mutant (S63A/S73A), T91 and T93 are still phosphorylated with significantly decreased rates when compared to the rates of S63 and S73 in the wild-type, despite being the only substrates of the kinase in this case (Fig. S3a & S4a). These data therefore suggest that the differences in the rates of the c-Jun N-terminal phosphorylation by JNK1 are determined primarily by the preference of the kinase for the primary sequences of the S-P rather than the T-P sites (Fig. 4c).

**Figure 4.**
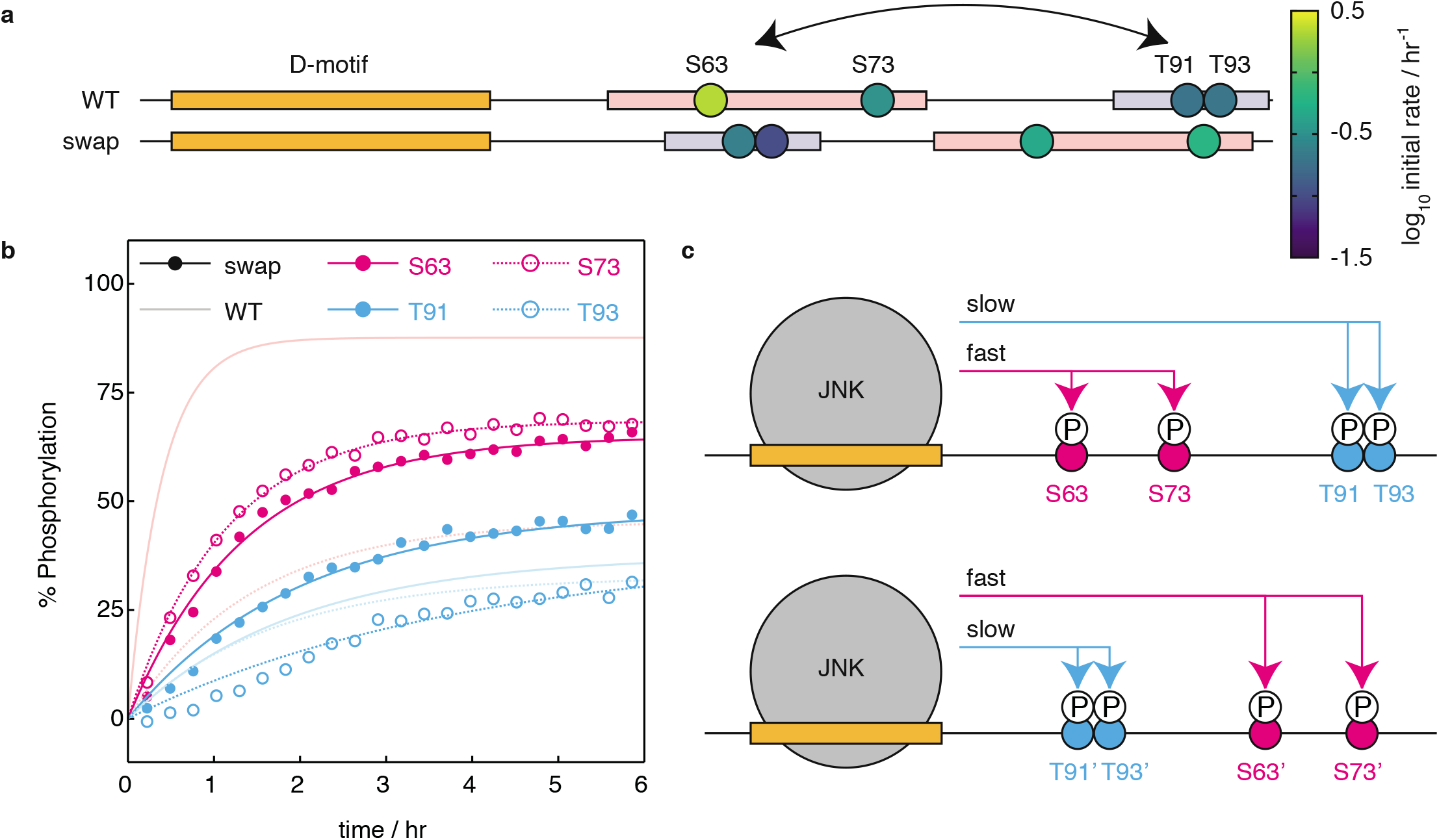
Effect of primary sequence of the S/T-P sites on the c-Jun N-terminal phosphorylation kinetics. (**a**) Schematic representation of the c-Jun swap TAD construct comparing to wild-type (WT). The initial rate of phosphorylation of residues by JNK1, determined by time-resolved NMR spectroscopy, is indicated according to the displayed colour scale. (**b**) Site-specific buildup curves of the phosphorylation of c-Jun swap mutant c-Jun TAD by JNK1; blurred lines show wild-type c-Jun TAD for comparison. (**c**) Schematic representation summarizing the effect of the primary sequences of the c-Jun TAD S/T-P residues on the c-Jun N-terminal progressive phosphorylation.

### The different kinetic groups of c-Jun N-terminal phosphorylation act positively and nagetively to regulate transcription by orchestrating the binding of co-regulators

Having identified groups of fast (S63, S73) and slow (T91, T93) phosphorylation sites, we next explored the biological function of the kinetics of the c-Jun phosphorylation by testing the effect of each phosphorylation kinetic group of the c-Jun TAD on its ability to activate transcription. For that purpose, we first analysed the effect of each individual c-Jun phosphorylation site with a *c-jun* luciferase reporter gene assay. Ectopic expression of c-Jun alanine point mutants (S63A, S73A, T91A, T93A) had no effect on the reporter activity (Fig. 5a). However, ectopic expression of c-Jun bearing alanine mutations in both S63 and S73 sites (2SA) greatly decreased the reporter activity when compared to the wild-type protein, which is consistent with previous observations^34^ Strikingly, alanine substitution of both the slower T91 and T93 sites (2TA) exhibited increased reporter activity comparing to the wild-type. Lastly, alanine substitution of all four phosphorylation sites (4A) showed decreased reporter activity relative to wild-type protein, similar to that seen with the 2SA variant (Fig. 5a). Taken together, these data suggest that the two phosphorylation kinetic groups of the c-Jun TAD have distinct activating and inhibitory effects on c-Jun transcriptional activity.

**Figure 5.**
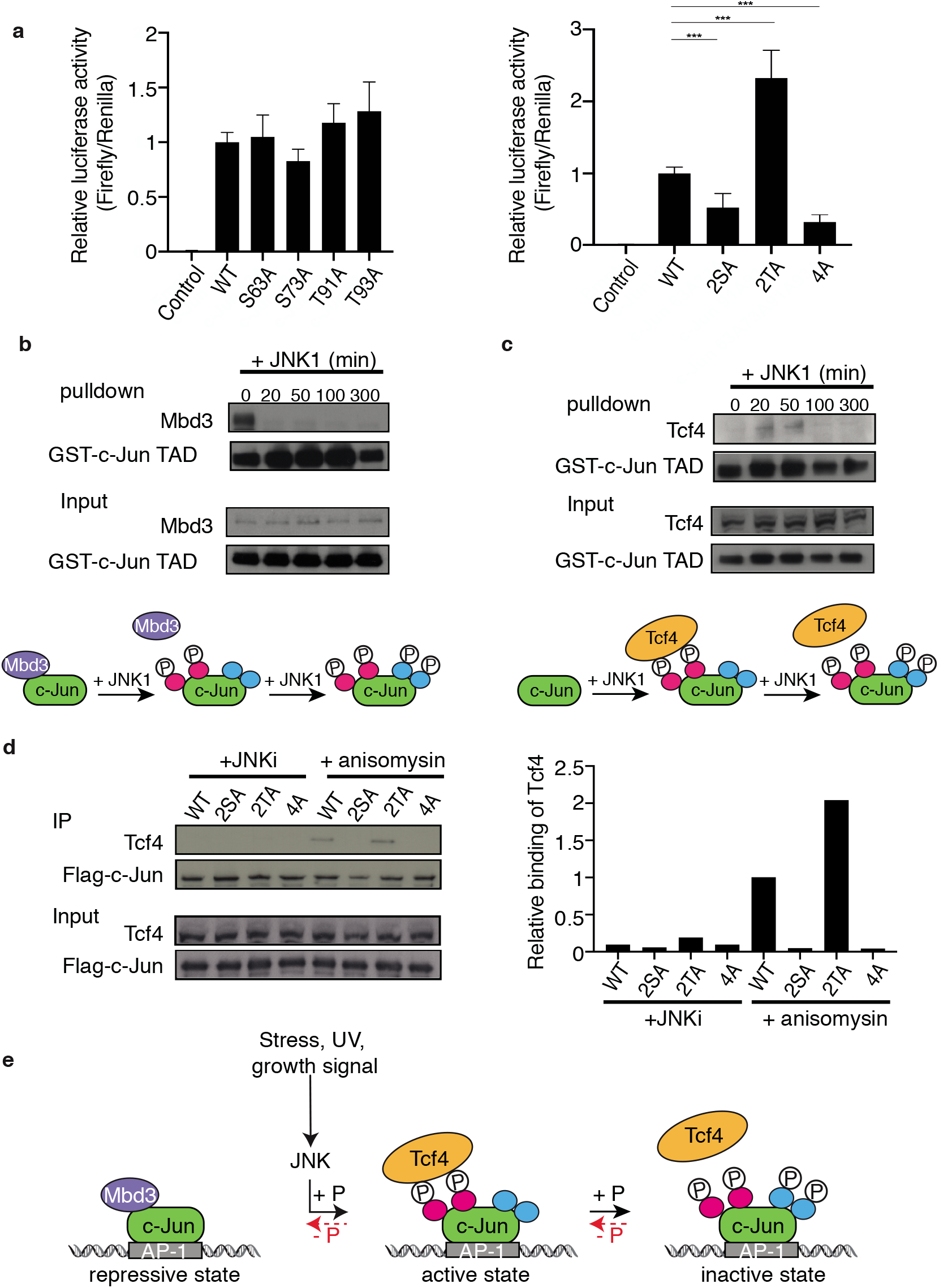
Effect of c-Jun temporal phosphorylation events on protein partners binding and transcriptional activity. (**a**) NIH-3T3 cells were transiently transfected to express: empty vector as a control (pFC2-dbd) or c-JunWT (pFA-c-JunWT) or c-Jun alanine mutants (Left; pFA-c-JunS63A, pFA-c-JunS73A, pFA-c-JunT91A or pFA-c-JunT93A. Right; pFA-c-Jun2SA, pFA-c-Jun2TA or pFA-c-Jun4A); firefly luciferase reporter gene (pFR-Luc); tk-Renilla as endogenous control (tk-Rluc); and constitutively active MEKK as an upstream activator of c-Jun phosphorylation (pFC-MEKK). Firefly luciferase activity was measured and normalized to Renilla luciferase activity and relative to c-JunWT. The graph represents the mean ± SD, calculated from at least three independent experiments performed in triplicate. * *p*<0.05, ** *p*<0.01 and *** *p*< 0.001 by Dunnett multiple comparison test. (**b and c**) HCT116 cell extracts were used in pulldown assays, using GST-c-Jun TAD as bait, phosphorylated by JNK1 at the indicated times. Recovered proteins were analysed by immunoblotting for (**b**) Mbd3 (**c**) Tcf4 (top). Schematic representation of the recovery of (**b**) Mbd3 and (**c**) Tcf4 from the performed pulldowns (bottom). (**d**) The indicated flag-tagged c-Jun wild-type or derivatives, transiently overexpressed in HCT116 cells, treated with either JNKi or stimulated additionally with anisomycin, were immunoprecipitated using Flag antibody. Immunoprecipitates were analysed for interaction with Tcf4 (left) and the quantification of the detected Tcf4 protein levels that were immunoprecipitated was performed using the Image Studio Lite Software (Licor), and normalized to the immunoprecipitated Flag-c-Jun and relative to WT + anisomycin (right). (**e**) Schematic summary model of the biological function of c-Jun N-terminal progressive phosphorylation by JNK. In cell resting conditions, c-Jun is in its unphosphorylated state and remains transcriptionally repressed by the recruitment of Mbd3. Upon JNK activation, c-Jun gets rapidly phosphorylated at S63 and S73 and Mbd3 exchanges with the Tcf4 transcriptional activator via its binding to the phosphorylated S63 and S73, activating transcription. As the signal progresses, JNK eventually phosphorylates T91 and T93, which inhibits Tcf4 binding and thus transcription. Eventually, secondary enzymes, such as phosphatases will restore c-Jun to its unphosphorylated repressive state that will rebind to Mbd3.

To gain further mechanistic insight on how the intrinsic kinetics of c-Jun N-terminal phosphorylation differentially control its transcriptional activity, we investigated the relationship between c-Jun phosphorylation and its interactions with the transcriptional repressor Mbd3 and the activator Tcf4. We conducted glutathione S-transferase (GST)-pull down experiments using recombinant GST-c-Jun TAD phosphorylated for different times such that phosphorylation was either partial or complete (Fig. 1d), and tested the recovery of endogenous Mbd3 and Tcf4 respectively from HCT116 cell extracts. In this assay, Mbd3 interacted exclusively with unphosphorylated c-Jun TAD, and binding was abolished as soon as c-Jun TAD phosphorylation was initiated (Fig. 5b). Interaction with Tcf4 was dependent on c-Jun TAD phosphorylation, and binding coincided with phosphorylation of S63 and S73, and was greatly reduced at the latest time point when c-Jun was fully phosphorylated at all four sites (Fig. 5c).

To examine the contribution of the different kinetic phosphorylation groups of c-Jun TAD to the Tcf4/phosphorylated c-Jun interaction, we performed co-immunoprecipitation assays using HCT-116 cells, transiently expressing Flag-tagged c-Jun or mutants (2SA, 2TA and 4A), to recover binding of endogenous Tcf4 from cell extracts treated either with JNK inhibitor (JNKi) or JNKi followed by anisomycin treatment to activate JNK (Fig. 5d). JNK phosphorylation and c-Jun phosphorylation of the individual S/T-P residues exhibit the same kinetics with anisomycin treatment in HCT116, as seen in NIH3T3 cells (Fig. S5a,b). This assay showed that Tcf4 binding is dependent on phosphorylation of S63 and S73, since their alanine substitution in both the 2SA and 4A c-Jun mutants did not allow recovery of Tcf4. Binding of Tcf4 was slightly enhanced in the case of the 2TA c-Jun TAD mutant compared to wild-type phosphorylated c-Jun TAD, indicating that T91/T93 phosphorylation may antagonise this interaction (Fig. 5d).

Taken together, and in agreement with the luciferase reporter gene assays (Fig. 5a), these data show that in unstimulated cells, c-Jun is not phosphorylated and is maintained transcriptionally repressed by binding to the Mbd3 subunit of the NurD repressor complex. Upon JNK activation, the Mbd3 transcriptional repressor exchanges with the Tcf4 transcriptional activator, via the rapidly phosphorylated S63 and S73 sites, thus triggering c-Jun transcriptional activity. However, sustained JNK signalling will lead eventually to phosphorylation of the remaining T91 and T93 sites, which abolishes the binding of Tcf4 and thus inhibits c-Jun activity.

## Conclusions

In this study we have explored the multisite phosphorylation of the c-Jun TAD by JNK, both *in vivo* and *in vitro*, and have identified a hierarchy of phosphorylation rates that leads to an intrinsic temporal signalling response. We have shown that c-Jun N-terminal phosphorylation is associated with three distinct functional phosphorylated states. Unphosphorylated c-Jun remains repressed via its binding to the Mdb3 subunit of the NurD repressor complex. Rapid phosphorylation of S63/S73 leads to an active state in which the Mdb3 repressor is displaced from the repressive, unphosphorylated state, allowing binding of the transcriptional activator Tcf4. Subsequently, slower phosphorylation of T91/T93 restores c-Jun to an inactive state in which Tcf4 binding becomes disfavoured. This behaviour thus provides a self-limiting safety mechanism to attenuate JNK signalling, independent of the engagement of phosphatases that must ultimately restore c-Jun to its repressive, unphosphorylated state (Fig. 5e).

The c-Jun TAD is now the second example of a transcription factor, after the TCF Elk-1^13^, for which an intrinsic hierarchy of phosphorylation rates has been shown to orchestrate the engagement of protein partners, leading to the autoregulation of a complex biological response elicited by a linear input signal. The existence of intrinsic functional kinetics of individual phosphorylation sites within multisite phosphorylated transcription factors thus appears to be a more general phenomenon, at least within the MAPK signalling pathway. For both c-Jun and Elk-1, phosphorylation kinetics convert the initial kinase signal into a pulse of transcriptional activity, however using different mechanisms. We have previously shown that the temporal kinetics of multisite phosphorylation of Elk-1 TAD controls the interaction with a single protein, Med23 subunit of the Mediator complex, to both positively and negatively regulate the transcriptional activity^13^. In contrast, the multisite phosphorylation of c-Jun orchestrates the coordinated recruitment and displacement of co-factor proteins, the combination of which determines the biological response. The response of c-Jun to phosphorylation is therefore potentially more complex, as the signalling output is dependent on interactions with several protein partners. For example, c-Jun phosphorylation is essential for intestinal tumor formation in epithelial cells via the Mbd3 and Tcf4 interactions^16,21^ yet triggers apoptosis in neuronal cells via its interaction with Bag1-L co-activator^19,35,36^. It appears likely that such a mechanism would allow binding of differential readers, e.g. tissue specific repressors and activators, of the c-Jun N-terminal phosphorylation kinetics to elicit distinct biological responses in different tissue types.

We have found that JNK kinase exhibits a strong preference for the primary sequences of the S63 and S73 sites within the c-Jun TAD (Fig. 3). The distance between the kinase binding site and the phosphorylation site, which modulates the effective concentration of the phosphorylation site^37^, did not strongly influence the observed phosphorylation kinetics. However, the relative orientation of docking and phosphorylation sites was critical, indicating that steric accessibility remains a significant factor. A sterically favoured positioning downstream of the D-motif rather than upstream, similarly to what we have observed for c-Jun TAD, has also been shown to be required for efficient phosphorylation of substrate sites by the MAP kinase ERK^38,39^.

A comparison of the mechanisms by which the MAP kinases JNK1 and ERK2 kinases recognise their substrate phosphorylation sites on c-Jun and Elk-1 TAD respectively, to generate the kinetics of multisite phosphorylation shows significant differences. The similar phosphorylation rates observed for residues in different c-Jun alanine variants provide insight into the mechanism of multisite phosphorylation by JNK1, indicating that the JNK1 active site cannot be completely saturated by the c-Jun phospho-sites, as the elimination of one phosphorylation site does not lead to a redistribution to other sites. This appears to contrast with the competitive mechanism of multisite phosphorylation observed in Elk-1, in which its active site is saturated by substrates such that elimination of a certain kinetic group of phosphosites results in a redistribution of the kinase to the remaining sites and hence more rapid phosphorylation^13^. This competitive model of Elk-1 multisite phosphorylation is governed by the position of individual Elk-1 substrate sites relative to the kinase docking interactions, and does not allow substrate site preference by the kinase, unlike what we observe for c-Jun. These differences could reflect the fact that Elk-1 is targeted by different MAPK kinases (ERK, p38 and JNK) each generating distinct phosphorylation kinetics^13^. Such a competitive mechanism appears more flexible, allowing for different phosphorylation patterns depending on the kinase. In the case of c-Jun, which is exclusively phosphorylated by JNK family kinases,, substrate site preference appears to be a more conservative mechanism that would not only ensure accurate timing of transcriptional activation and subsequent elimination but potentially filter binding of other MAPK kinases^40^.

The identification of three functional states associated with c-Jun N-terminal phosphorylation may have implications for the development of novel therapeutic cancer strategies. The JNK signalling pathway has been a desirable drug target for years, but so far JNK inhibitors have not been translated into clinical use. The main reason is the lack of specificity of the current inhibitors and cellular toxicity^41,42^. JNK plays an important role in various normal biological functions and its direct inhibition could have undesired side effects. Inhibitors targeting specific JNK-mediated downstream substrates and cellular events, such as phospho-c-Jun or the interaction of phospho-c-Jun binding partners, may show increased tumour specificity and efficacy. Our results suggest that in the case of c-Jun, the active double phosphorylated pS63/pS73 c-Jun form, and its interaction with activators, would be the most promising candidates for more personalised therapeutic intervention.

In summary, in this study we have shown that c-Jun multisite phosphorylation by JNK exhibits an intrinsic kinetics that leads to a precise and timed regulation of the transcriptional output of the JNK signalling pathway, where both positive and negative outputs are generated by a single kinase input. The concept of the temporal phosphorylation pattern and its translation to a self-eliminating transcriptional mechanism thus appear to be more general in the MAPK signalling. This mechanism contrasts the dogma of the classic ‘on and off switch’ of the transcriptional response via the dual role of kinases and phosphatases in MAPK signalling and perhaps more general in signalling pathways, revealing that the temporal dimension of multisite phosphorylation of a protein domain allows for more elaborate signalling responses. We speculate that the intrinsic kinetics of post-translational modifications across multiple sites might provide a simple and universal mechanism for the generation of a complex temporal response to an initial signalling event, and represent a generalisation of the concept of proteins exhibiting posttranslational modifications at multiple sites that potentially serves to regulate any type of complex biological response.

## Methods

### Molecular techniques

Plasmid constructions, protein analysis by SDS-PAGE and immunoblotting used standard methods. c-Jun point mutations were generated by PCR-based mutagenesis using the site-directed mutagenesis kit (New England Biolabs). cDock, mDock and swap c-Jun constructs were chemically synthesized by ATCC.

### Antibodies

pS63 c-Jun (9261), pS73 c-Jun (9164), JNK (9252) and p183p185 JNK (9251) from Cell Signalling; pT91 c-Jun (ab247509), and pT93 c-Jun (ab81319) from Abcam; Mbd3 (A302-529A) from Bethyl Laboratories; c-Jun (554083) from BD biosciences; Tcf4 (PA5-25659) from Thermo Fisher; GST (G7781) and Flag (F3165) from Sigma-Aldrich.

### Expression and purification of recombinant GST-c-Jun TAD

Wild-type or mutant human c-Jun sequences encoding residues 1-151 were inserted between the BamHI and NotI sites of modified pET-41a^13^ and expressed as GST-(His)6 fusion proteins. Protein expression was induced in *E. coli* Rosetta(DE3) (Novagen) with 0.5 mM IPTG for 3h at 37°C. Cells were lysed in 50 mM Tris pH 8.0, 300 mM NaCl, 5 mM DTT, with complete protease inhibitor cocktail (Merck). Cleared lysate was applied to glutathione sepharose resin (GE Healthcare Life Sciences), incubated for 1h with the resin, and then washed extensively with lysis buffer. Proteins were eluted with 20mM reduced glutathione in 50 mM Tris pH 8.0, 150 mM NaCl, 5mM DTT, and further purified on a Superdex 200 HR10/30 size exclusion column (GE Healthcare Life Sciences) in 10 mM K_2_HPO_4_ pH 6.8, 50 mM NaCl. For^15^N-and^15^N-/^13^C-labelling, cells were grown in M9 minimal medium containing^15^NH_4_Cl and^13^C-glucose as required and c-Jun was expressed for 3h at 37°C after induction with 0.5 mM IPTG. After adsorption to glutathione sepharose the c-Jun TAD was released by incubation with 1:50 (w/w) GST-3C protease on the resin overnight at 4°C, and purified by size exclusion chromatography as above.

### NMR spectroscopy

NMR data for assignment of c-Jun backbone resonances were acquired at 283 K using a Bruker Avance III NMR spectrometer with TXI cryoprobe operating at 700 MHz (^1^H Larmor frequency). A 150 μM sample of uniformly^15^N,^13^C-labelled sample of c-Jun was prepared in 10 mM K_2_HPO_4_ pH 6.8, 150 mM NaCl, 5 mM MgCl_2_, 10% (v/v) D_2_O and 0.01% (w/v) DSS. BEST-HNCO, HNCACO, HNCACB and HNCOCACB experiments^43^ were acquired, processed using nmrPipe^44^ and analysed using CCPN Analysis^45^. Secondary structure populations were calculated from HN, N, CO, CA and CB chemical shifts using δ2D^28^. c-Jun phosphorylation kinetics were measured at 20°C using a Bruker Avance III NMR spectrometer with TXI cryoprobe operating at 950 MHz (^1^H Larmor frequency). 391 μL samples of^15^N-labelled cJun (WT or variants) were prepared in Shigemi tubes without plungers, at concentrations of ca. 75 μM in 10 mM K_2_HPO_4_ pH 6.8, 50 mM NaCl, 5 mM MgCl_2_, 2 mM DTT, 1 mM ATP, 5% (v/v) D_2_O. Reference 2D^1^H,^15^N SOFAST-HMQC experiments^46^ were acquired using a 50 ms recycle delay, with 1536 complex points and a sweep width of 16 ppm in the direct dimension (100 ms acquisition time), and 64 complex points and a sweep width of 20 ppm in the indirect dimension (33.2 ms acquisition time). Phosphorylation was then initiated by rapid mixing with 290 U recombinant active JNK1 (Proqinase), and a series of SOFAST-HMQC experiments were acquired over a period of several hours, with 4 scans per increment (corresponding to 93 seconds per spectrum). Spectra were processed in nmrPipe^44^ using linear prediction and cosine-squared window functions, then imported into Julia or analysis with NMRTools.jl. Following previous studies^13,23,24^ box integrals were calculated around phosphorylated resonances and fitted to single exponential functions as a function of time.

### Cell Culture

NIH3T3 and HCT116 were cultured in Dulbeccos’s Modified Eagle Medium (DMEM) supplemented with 10% fetal bovine serum (FBS). Cells were treated when indicated with 50 μM JNK inhibitor (JNKi) (SP600125, Calbiochem) or 15 ng/ml anisomycin (Sigma). HCT116 cells were plated at subconfluence and transfected with lipofectamine 2000 (Invitrogen) when indicated.

### Phosphorylation assays

For phosphorylation of GST-c-Jun TAD, or the^15^N isotope-labeled protein version, 75 μM of substrate was incubated with 290 U recombinant active JNK1 (Proqinase) in 10 mM K_2_HPO_4_ pH 6.8, 50 mM NaCl, 1 mM ATP, 2mM DTT, 5 mM MgCl_2_ at 20°C for the indicated times.

For phosphorylation of endogenous JNK and c-Jun, cells were treated with 50 μM of JNKi for 2h followed by 15 ng/ml anisomycin treatment for the indicated times at 37°C. Lysates for SDS-PAGE were prepared in 300 μl RIPA buffer (Thermo Fisher) supplemented with protease inhibitors (Sigma), 1 mM NaF and 1 mM Na_3_VO_4_. Protein quantification was performed with the BCA kit (Thermo Scientific) and densitometry analysis of the immunoblot with the Image Studio Lite Software (Licor).

### Reporter gene assay

NIH 3T3 cells were transiently transfected with pFA2-cJunWT (PathDetect c-Jun *trans-Reporting* System from Agilent Technologies), or the indicated pFA2-cJun mutants or pFC2-dbd (empty vector); pFR-Luc; pFC-MEKK; and tk-Renilla, using Lipofectamine 2000 (Invitrogen). Next day, medium was replaced with starvation medium (0.3% FBS) for 24 h. Luminescence activity was measured 48h post-transfection using Dual-Luciferase Reporter Assay System following manufacturer’s protocol (Promega) in a FLUOStar Omega Plate reader (BMG Labtech). Data were expressed as fold induction after being normalized using tk-renilla luciferase. The data are presented as the mean ± standard deviation (SD), and compared using Dunnett multiple comparisons test using GraphPad Prism version 8.0 (Graphpad Software Inc.). p values less than 0.05 were considered statistically significant (p<0.05 *; p<0.01 **; p< 0.005 ***).

### Immunoprecipitation

HCT116 cells were transiently transfected with pCMV-Tag2-Flag-cJun (provided by Axel Behrens) wild-type or mutants, and were treated with JNKi or additionaly with anisomycin. Cells were lysed in Lysis Buffer (PBS with 0.1% NP-40) with protease inhibitor (Merck), 1 mM Na_3_VO_4_, 1 mM NaF, and 1 mM PMSF, sonicated and cleared by centrifugation for 15 min at 13,000 rpm at 4 °C. Lysates were incubated with 20 μl ANTI-FLAG M2 affinity gel (Sigma) for 3 h at 4 °C. Beads were washed 3 times with Lysis Buffer (PBS with 0.1% NP-40) and proteins were eluted with 20 μl of 1x SDS loading buffer.

### GST-fusion affinity assays

HCT116 cells were lysed in RIPA buffer (for Mbd3) or Lysis Buffer (PBS with 0.1% NP-40) (for Tcf4) with protease inhibitor cocktail (Merck), 1 mM Na_3_VO_4_, 1 mM NaF, and 1 mM PMSF, sonicated and cleared by centrifugation for 15 min at 13,000 rpm at 4 °C. 10 μl of gluthatione sepharose beads (GE Healthcare) were incubated with 10 μg GST c-Jun TAD fusion protein for 1h at 4 °C. After binding beads were washed 3 times with binding buffer (PBS with protease inhibitor cocktail (Merck) and 1 mM Na_3_VO_4_) and were next incubated with the lysed proteins for 3 h at 4 °C. Beads were washed 3 times with with Wash Buffer (50 mM Tris pH 7.5, 250 mM NaCl, 0.5% NP40, 5 mM EDTA and 1 mM DTT) (for Mbd3) or Lysis Buffer (PBS with 0.1% NP-40) (for Tcf4) and elution step was performed the same way as for the immunoprecipitation assay (see above).

## Acknowledgements

We thank Gabriel Waksman and Carolyn Moores for support and fruitful discussions. Natalie Barthel and Jordi Gines for technical support with biochemical and NMR experiments. Alkistis Mitropoulou and Lisa Cabrita for help and advice with labelling of recombinant c-Jun TAD, Angelo Figueiredo for help with initial NMR measurements. Clive Da Costa and members of the Behrens lab for a lot of support and technical advice. We also thank Claire Bagneris for lab management assistance. AM was supported by a Wellcome / Birkbeck ISSF Early Career Fellowship.

The authors have no conflict of interests.

## Author contributions

S.A-T., S.S., P.R.B and A.M. designed and performed all biochemical, molecular and cell biology experiments; C.W. designed, executed, and interpreted the NMR data analysis experiments together with J.C.; A.B provided reagents and interpreted the experiments. A.M. conceived the project, designed, interpreted and supervised all experiments; A.M. wrote the paper with C.W and S.A-T and A.B..

