## Supplementary figures and legends for "An intrinsic temporal order of c-Jun N-terminal phosphorylation regulates its activity by orchestrating co-factor recruitment"

**Fig. S1.** (**a**) $^{1}H,^{15}N$ HSQC spectrum of the c-Jun TAD (283 K, 700 MHz), showing resonance assignments. Assignments marked with an asterisk indicate folded resonances. (**b**) Secondary structure populations within the c-Jun TAD determined from HN, N, C', CA and CB chemical shifts using δ2D [28].

**S1**

**a**

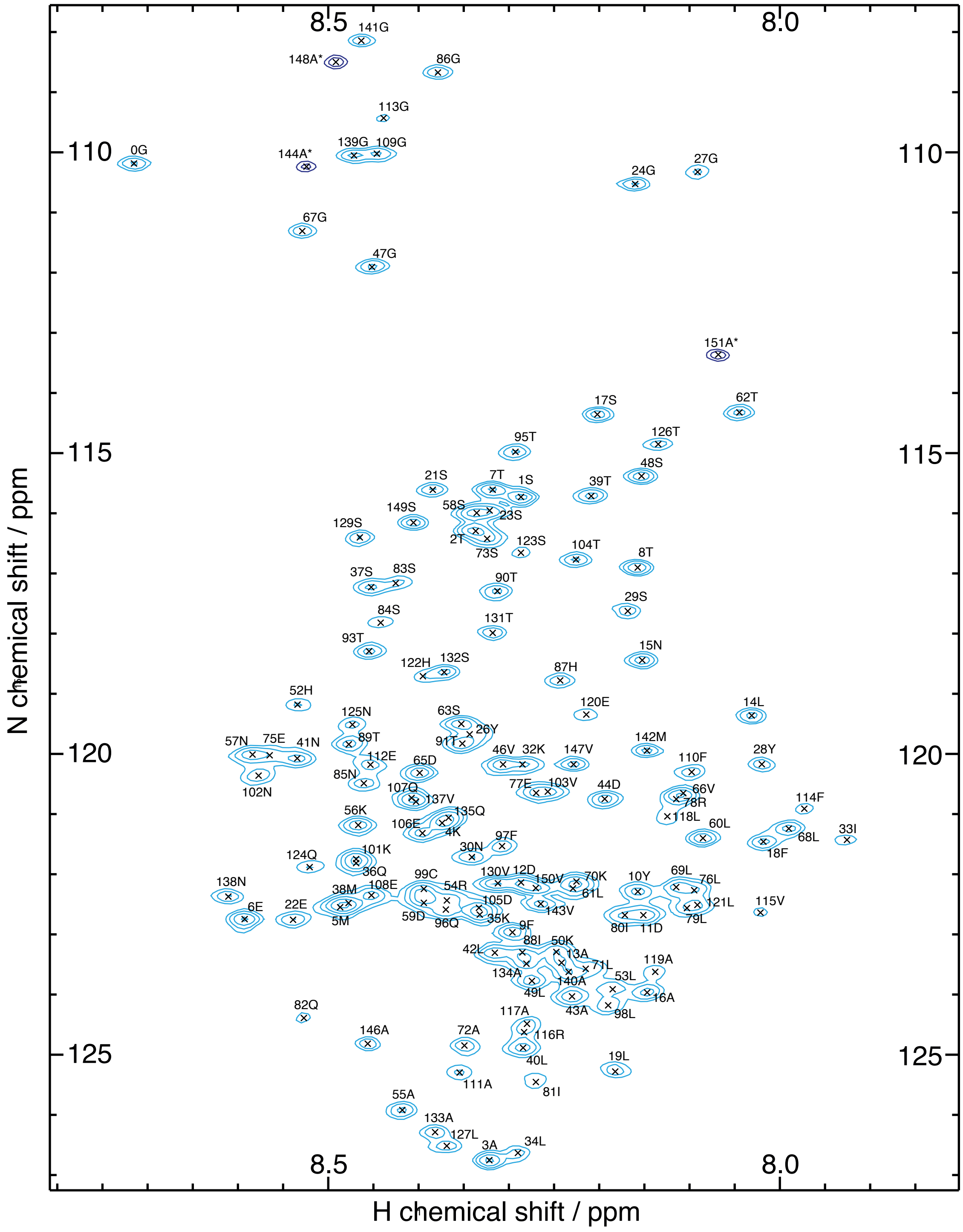

8.5
8.0
110
115
120
125
N chemical shift / ppm
H chemical shift / ppm
141G
148A*
86G
113G
0G
144A*
139G
109G
24G
27G
67G
47G
151A*
17S
62T
95T
126T
21S
7T
1S
48S
39T
149S
58S
23S
129S
2T
73S
123S
104T
8T
37S
83S
90T
29S
84S
131T
93T
15N
122H
132S
87H
52H
120E
14L
125N
63S
26Y
57N
75E
41N
89T
91T
46V
32K
147V
142M
28Y
110F
112E
65D
102N
85N
77E
103V
107Q
44D
66V
137V
78R
114F
118L
56K
135Q
106E
4K
60L
33I
68L
97F
18F
30N
101K
124Q
36Q
130V
12D
70K
10Y
69L
99C
150V
76L
138N
38M
108E
54R
81L
61L
105D
121L
115V
6E
22E
5M
59D
96Q
35K
143V
80I
11D
79L
9F
88I
50K
42L
13A
71L
119A
134A
140A
53L
49L
16A
43A
98L
82Q
117A
146A
72A
116R
40L
19L
111A
81I
55A
133A
127L
3A
34L


**b**

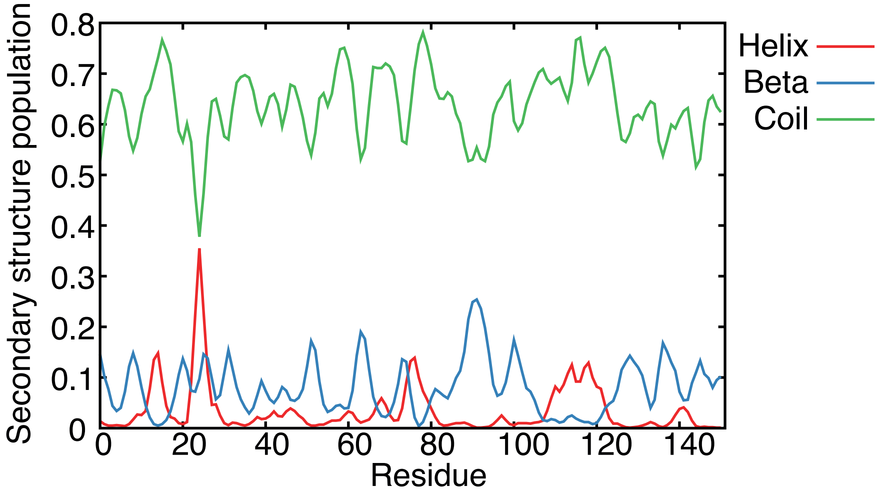

Secondary structure population
0.8
0.7
0.6
0.5
0.4
0.3
0.2
0.1
0
0
20
40
60
80
100
120
140
Residue
Helix
Beta
Coil

**Fig. S2.** $^{1}H,^{15}N$ SOFAST-HMQC spectra of WT c-Jun (293 K, 950 MHz) acquired at the indicated times following addition of active JNK1 to initiate phosphorylation. A spectrum of unphosphorylated c-Jun is shown in grey for reference. Phosphorylated resonances have been highlighted in pink and cyan according to the kinetic group (fast / slow) with assignments as indicated. We note that two well-separated resonances were observed for pS63. In all analyses, we use the sum of these resonances as a reporter of pS63.

**S2**

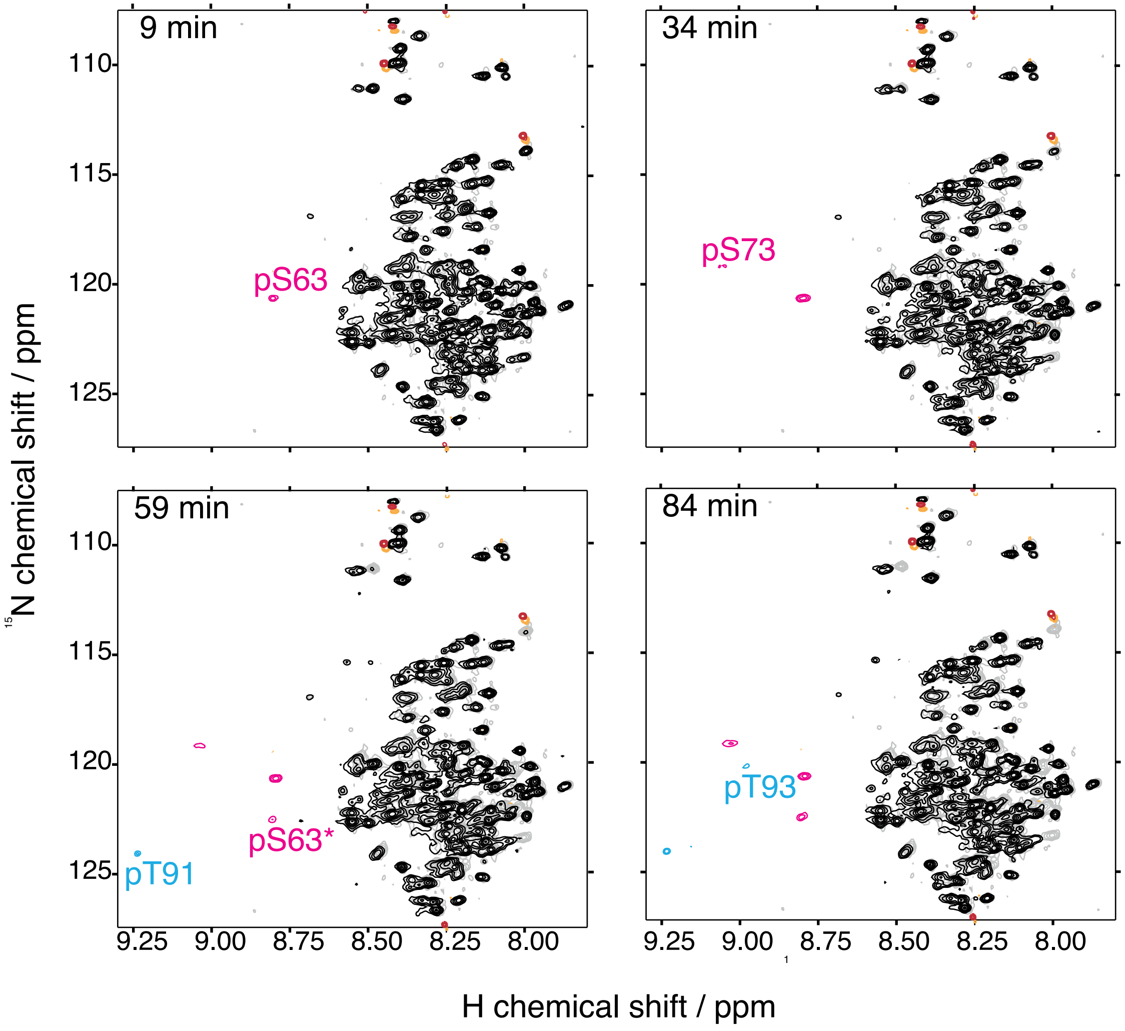
9 min
34 min
59 min
84 min
pS63
pS73
pS63*
pT91
pT93
110
115
120
125
9.25
9.00
8.75
8.50
8.25
8.00
15N chemical shift / ppm
1H chemical shift / ppm

**Fig. S3.** (**a**) Assignment of phosphorylated c-Jun amide resonances. Contour plots indicate the maximum values observed in $^{1}H,^{15}N$ SOFAST-HMQC spectra (293 K, 950 MHz) across a phosphorylation time course, for wild-type (WT) c-Jun (grey) and variants as indicated (red). Resonance assignments are indicated on each panel. (**b**) Phospho-specific immunoblotting analysis of recombinant wild-type and alanine point mutants (S63A, S73A, T91, T93A) c-Jun TAD *in vitro*. c-Jun TAD proteins were purified and phosphorylated by JNK1 for 400 min as described in Materials and Methods section.

**S3**

**a**

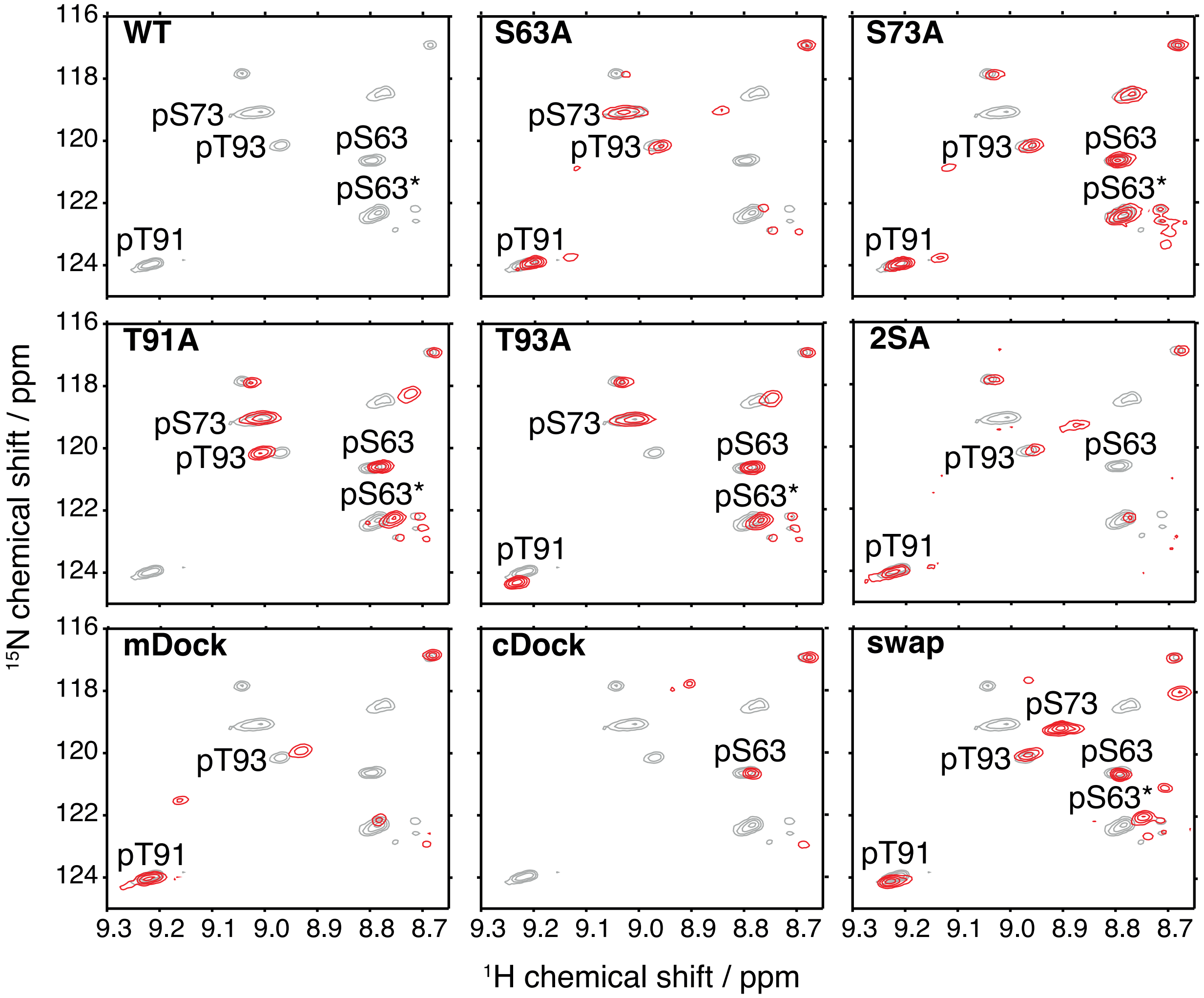
WT
S63A
S73A
T91A
T93A
2SA
mDock
cDock
swap
pS73
pT93
pS63
pS63*
pT91
116
118
120
122
124
9.3 9.2 9.1 9.0 8.9 8.8 8.7
$^{15}$N chemical shift / ppm
$^{1}$H chemical shift / ppm

**b**

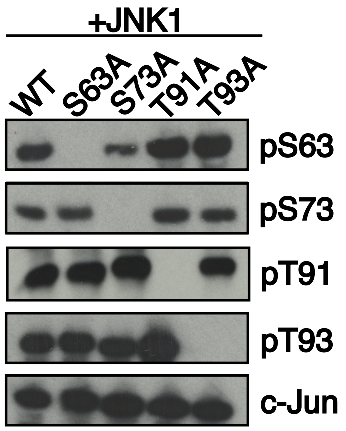
+JNK1
WT
S63A
S73A
T91A
T93A
pS63
pS73
pT91
pT93
c-Jun

**Fig. S4.** (**a**) Phosphorylation time-courses obtained by integration of phosphorylated resonances indicated in the legend, for c-Jun variants (S63A, S73A, T91A, T93A and 2SA) comparing to wild-type (WT). Measurements are fitted to single exponential build-up curves. (**b**) Amino acid sequence changes of the mDock and cDock c-Jun TAD constructs analysed by time-resolved NMR, comparing to the WT sequence. (**c**) Amino acid sequence changes of the swap c-Jun TAD construct analysed by time-resolved NMR, comparing to the WT sequence.

**S4**

**a**

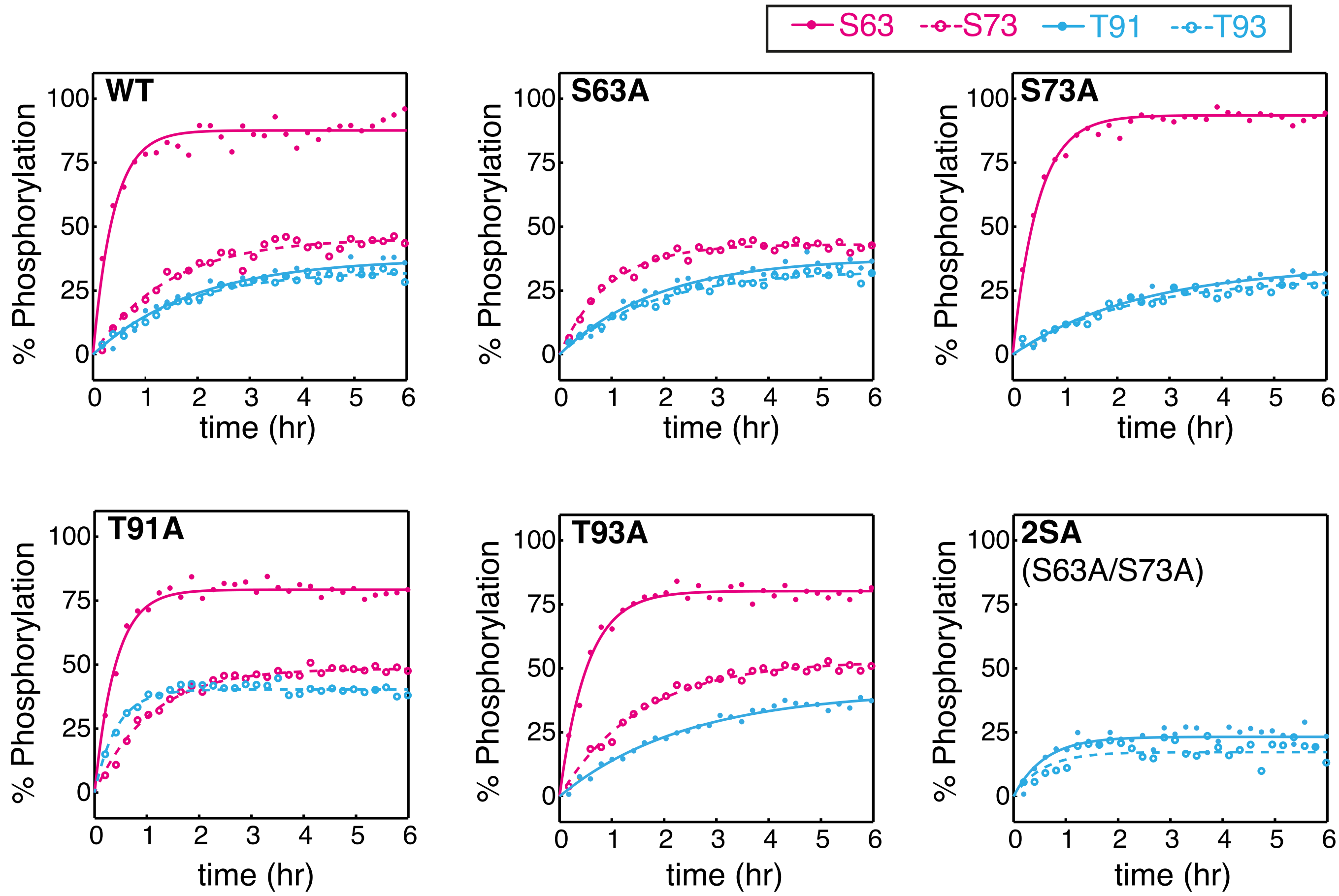


**b**

**WT c-Jun TAD** 1-MTAKMETTFYDDALNASFLPSESGPYGYSNPKILKQSMTLNLADPVGSLKPHLRA
KNSDLLTSPDVGLLKLASPELERLIIQSSNGHITTTPTPTQFLCPKNVTDEQEGF
AEGFVRALAELHSQNTLPSVTSAAQPVNGAGMVAPAVASVA-151

**mDOCK c-Jun TAD** 1-MTAKMETTFYDDALNASFLPSESGPYGYSNPPHLRAKNSDLLTSPDVGLLKLASP
ELERLIIQKILKQSMTLNLADPVGSLKSSNGHITTTPTPTQFLCPKNVTDEQEGF
AEGFVRALAELHSQNTLPSVTSAAQPVNGAGMVAPAVASVA-151

**cDOCK c-Jun TAD** 1-MTAKMETTFYDDALNASFLPSESGPYGYSNPPHLRAKNSDLLTSPDVGLLKLASP
ELERLIIQSSNGHITTTPTPTQFLCPKNKILKQSMTLNLADPVGSLKVTDEQEGF
AEGFVRALAELHSQNTLPSVTSAAQPVNGAGMVAPAVASVA-151

**c**

63 73 91 93

**WT c-Jun TAD** N-NSDLLTSPDVGLLKLASPELERL-/-NGHITTTPTPTQFLC-C

91 93 63 73

**swap c-Jun TAD** N-NGHITTTPTPTQFLC-/-NSDLLTSPDVGLLKLASPELERL-C

**Fig. S5** (**a**) Time-course immunoblot analysis of endogenous JNK phosphorylation/activation in HCT116 cells following anisomycin treatment for the indicated times (top), and quantification of the detected protein levels by western blot using Image Studio Lite Software (Licor) and normalized to total JNK expression (bottom). (**b**) Phospho-specific immunoblotting analysis of endogenous c-Jun time-course phosphorylation in HCT116 cells following anisomycin treatment for the indicated times (top), and quantification of the detected protein levels by western blot using Image Studio Lite Software (Licor) normalized to total c-Jun expression (bottom).

**S5**

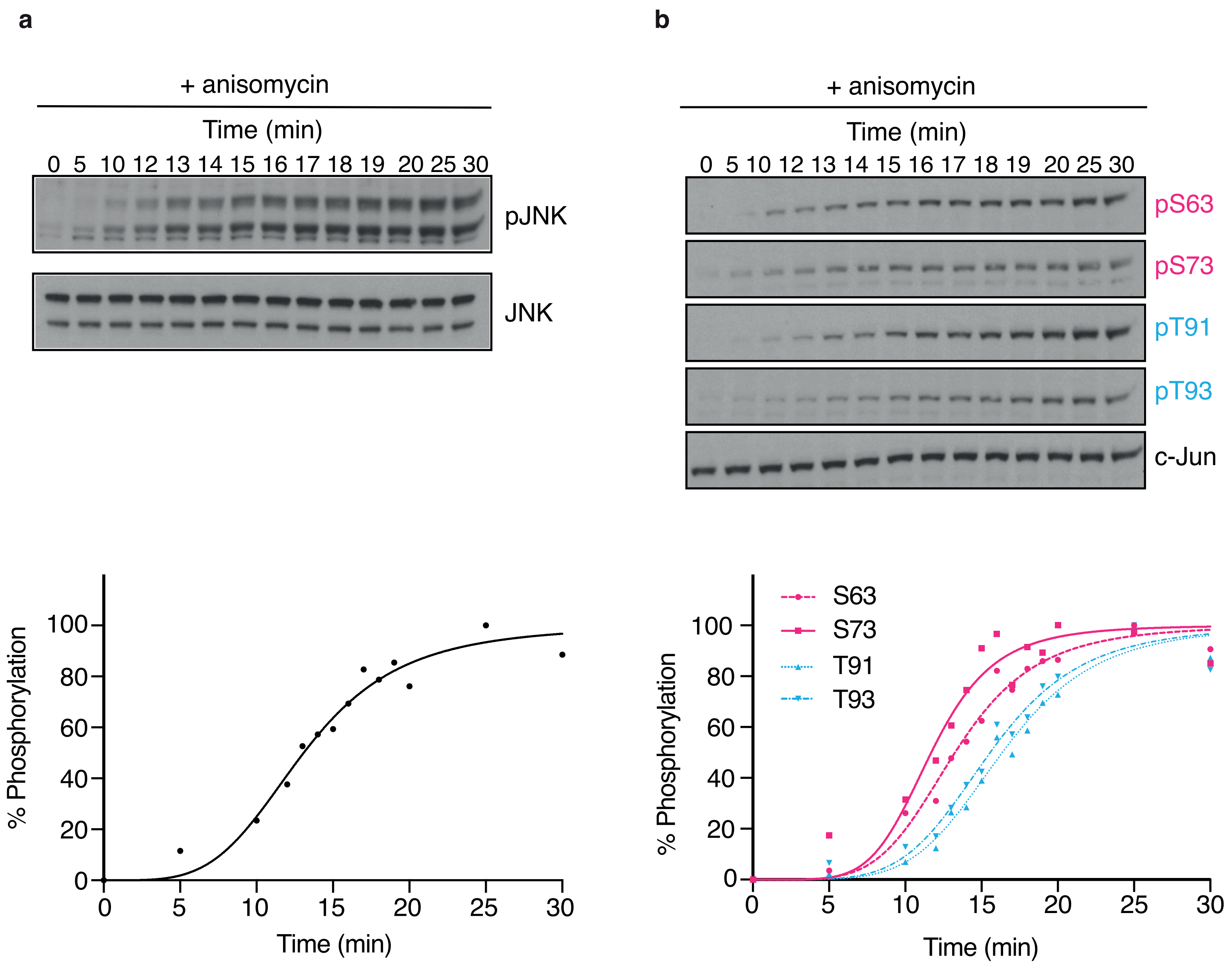
a
+ anisomycin
Time (min)
0 5 10 12 13 14 15 16 17 18 19 20 25 30
pJNK
JNK
% Phosphorylation
0
20
40
60
80
100
0 5 10 15 20 25 30
Time (min)
b
+ anisomycin
Time (min)
0 5 10 12 13 14 15 16 17 18 19 20 25 30
pS63
pS73
pT91
pT93
c-Jun
S63
S73
T91
T93
% Phosphorylation
Time (min)
